# Comparative analysis of HPV-Human Protein Interaction Networks in Oropharyngeal and Oral Squamous cell carcinomas

**DOI:** 10.1101/2023.12.05.570064

**Authors:** Arsalan Riaz, Faisal F. Khan

## Abstract

Oral Squamous Cell Carcinoma (OSCC) accounts for more than 90% of reported head and neck cancers which is the sixth leading cancer worldwide. Oropharyngeal squamous cell carcinoma (OPSCC) cases, a closely located but a different sub-type of head and neck cancer, are also rising worldwide specifically among young individuals with risk factors including viral infection, smoking and alcohol. HPV infection specifically the subtype HPV16 is a significant risk factor in the initiation and progression of OPSCC, but has not been reported to play a role in OSCC. This study sets out to decipher mechanistic differences at the molecular level by comparing the viral-human protein-protein interaction (PPI) networks both in oropharyngeal squamous cell carcinoma (OPSCC) and oral squamous cell carcinoma (OSCC). We found that the unbiased HPV16-Human network consisted of 479 nodes, including 7 HPV16 proteins and 472 human proteins. Enrichment analysis revealed significant involvement of the identified human proteins in ubiquitination, protein degradation, and cancer-related pathways. A subset of 37 genes showed differential expression in HPV-positive OPSCC and OSCC, with TP53 over-expressed and ENDOD1 under-expressed in both cancers. Six genes (NUTM1, MYC, SCN9A, COL27A1, ITGB4, and GNB2) exhibited distinct changes in HPV-positive OPSCC compared to the other groups, where NUTM1 was the most over-expressed in OPSCC HPV+. The identified genes and pathways could serve as potential targets for precision medicine and therapeutic interventions in HPV-associated cancers. Further investigations are required to validate their clinical implication in *in vitro* and *in vivo* models.

## Introduction

Oral cancer is the sixth most common cancer around the world and fourth amongst the South Asian regions, mostly affecting middle and low-income countries. Oral Squamous Cell Carcinoma is the more prevalent type of oral cancer that targets a major chunk of the young population [1] but in current statistics, the incidence of Oropharyngeal Squamous Cell Carcinoma (OPSCC) is increasing as compared to other types of head and neck cancers including oral cancer [2, 3]. Risk factors for oral cancer include smoking of tobacco, alcohol consumption, unhygienic oral condition, excessive use of hookah (also *hukka*) and betel quid (also *gutka*), low fruit and vegetable consumption and viral infection especially the Human Papilloma Virus (HPV) [4]. HPV is classified into high-risk and low-risk types. Among the high-risk types, HPV 16 and 18 are most commonly involved in invasive cancer. HPV16 has been shown to cause cancer at the base of the tongue and tonsillar area in young individuals [5]. The Low-risk types, that present a low oncogenic risk, include HPV 6, 11, 40, 42, and a few more.

HPV is a non-enveloped, circular double-stranded DNA virus where the genome consists of 3 regions: Late (L) Region, Early (E) Region, and Long Control Region (LCR). The early region encodes 6 proteins which include E1, E2, E4, E5, E6, and E7 while L1 and L2 are expressed by the late region. Viral proteins are encoded by L and E regions while LCR contains the origin of viral DNA replication and transcriptional regulatory elements. HPV genes E6 and E7 which produce E6 and E7 oncoproteins confer the virus with oncogenic properties through p53 and retinoblastoma (Rb) inhibition. E6 causes ubiquitin-mediated proteolysis and degradation of p53 leading to increased mitotic stress and genome instability. On the other hand, E7 binds and inactivates tumour suppressor proteins of the retinoblastoma family which leads to abnormal cell proliferation and over-expression of tumour suppressor protein p16 [6].

HPV is regarded as a significant factor in Head and Neck Cancer particularly in oropharyngeal tumours, as compared to alcohol and tobacco [7]. HPV-positive oropharyngeal squamous cell carcinoma (OPSCC) represents different clinical and anatomical features compared to Oral Squamous Cell Carcinoma (OSCC) distinguished by the presence of tonsillar tissue, which also contains an actively dividing basal layer [8]. Previously, carcinoma developing from the oropharynx and oral cavity was considered to be a monotonous tumour because of the continuous squamous epithelium and was termed oral cancer as a whole [2] until it was updated in the 4th Edition of the World Health Organisation Classification of Head and Neck Tumours in 2017 [9]. The current state of knowledge shows the prevalence of HPV more towards OPSCC as compared to OSCC but the need for further exploration remains. Woods *et al*, suggest that other head and neck cancers are 5 times less likely to be attributed to HPV infection when compared with OPSCC [10].

The anatomical characteristics of HPV-positive OPSCC are established and the underlying mechanism of the viral deposition has also been studied [11], however, there is a gap in understanding the molecular signatures exclusive to HPV-positive OPSCC. Identifying these signatures can lead to new avenues in precision medicine, particularly for HPV16-induced OPSCC.

Protein-protein interaction networks can provide essential information for understanding the underlying molecular mechanisms of complex diseases, such as cancer [12]. During the course of a viral infection, protein-protein interactions between the viral and host proteins can highlight the proteins and associated pathways utilised by the virus to escape the host immune systems for replication and survival [13]. For this purpose, knowledge of HPV16-encoded protein interactions and the differential gene expression of the host proteins in infected cells can be important for gaining a deep understanding of associated pathways, especially in the context of OPSCC when compared with OSCC.

In this study, we create and analyse the core viral-host interaction network in OPSCC and OSCC and their associated pathways through network analysis. We further prioritised interesting candidate genes based on expression profiles in HPV-positive OPSCC patients.

## Methods

### Construction of HPV16-Human PPI network

Protein interaction data for Human Papillomavirus 16 was received form BioGrid (Release 4.4.214), and filtered for HPV16 (Taxon ID: 333760) and Human proteins (Taxon ID: 9606). The interaction data was then filtered for ‘physical’ interactions only, which were then used for network construction. The resulting network was visualised using Cytoscape (version 3.9.0). Self-loops and duplicate edges were removed while ignoring the edge direction. Finally, we used Network Analyzer (version 4.4.8) to analyse the network topology and compute summary statistics of the network.

### Enrichment analysis

We analyzed gene lists of interest using EnrichR, a web-based interface (https://maayanlab.cloud/Enrichr/), to conduct enrichment analysis of GO annotation and KEGG pathway terms. We used Entrez gene symbols as identifiers for the gene lists, and our gene-set libraries included KEGG 2021 Human, GO Biological Process 2021, GO Molecular Function 2021, and GO Cellular Component 2021. We utilised default parameters for the analysis and obtained a table of results that included p-value, adjusted P-Value, Odd Ratio, Combined Score, and overlapping genes. The overlapping genes represented the number of genes in the input gene list that overlapped with the genes in the enriched term or pathway. EnrichR computes the p-value using the fisher exact test, which assumes a binomial distribution and independence for the probability of any gene belonging to any set. The adjusted p-value is computed using the Benjamini-Hochberg method for correction for multiple hypotheses testing. The combined score is derived from running the Fisher’s exact test for many random genes sets to compute a mean rank and standard deviation from the expected rank for each term in the gene-set library and finally calculating a z-score to assess the deviation from the expected rank [14-16]. The terms with P-Value greater than 0.05 were considered insignificant.

### Retrieval of gene lists with mutations reported in OSCC and OPSCC

Gene lists with mutations reported in OSCC and OPSCC were retrieved from cBioPortal (v3.5.4) for cancer genomics [17]. The Cancer Genome Atlas studies were considered, and samples with known HPV status of the primary oral cavity and oropharynx tumours were considered. For the oropharynx, only 33 samples were available of which 22 were HPV+ and 11 were HPV-, for the oral cavity 196 samples were available of which only 12 were HPV+ and 160 were HPV-. The frequency cut-off was set to 2.6% (total genes = 9796) for the oral cavity and for the oropharynx the frequency cut-off was set to 6.1% (total genes = 3043), due to an insignificant number of samples, oropharynx additional genes were identified via literature review on HPV associated OPSCC. The literature review identified mutations in 73 additional unique genes that were reported in OPSCC samples and were added to the original mutation list, resulting in a final list of mutations in 560 genes for OPSCC and 612 genes for OSCC (Supplementary Table ST1 and ST2).

### Differential gene expression analysis

Raw counts data was retrieved from the GDAC firehose (release 2016_01_28) (https://gdac.broadinstitute.org/) hosted by the Broad Institute. We compared primary tumour cases of oropharynx both HPV positive (n=22) and negative (n=10) and OSCC both HPV positive (n=11) and negative (n=142) against normal (n=31) for differential expression analysis using R package DESeq2 (version 1.38.2), which implements a model based on the negative binomial distribution [18].

## Results

### Understanding the Viral-Host Protein-Protein Interaction Network

We built a protein-protein interaction (PPI) network between human proteins and HPV16 proteins, with the interaction data coming from BioGRID. After removing self-loops and duplicate edges, we analysed the connectivity of 479 nodes, 7 of which were HPV16 proteins and 472 of which were human proteins, having 506 edges in the PPI. E5, E6, and E7 were the top hub proteins with a high degree of interaction (Figure 1). No densely connected nodes were present, as indicated by the clustering coefficient was 0 (Table 1). Furthermore, the PPI network’s high heterogeneity and low density demonstrate its scale-free nature, as it indicates that there are a few highly connected hub proteins that interact with many other proteins, while the majority of proteins have relatively few connections [19].

**Figure 1:**
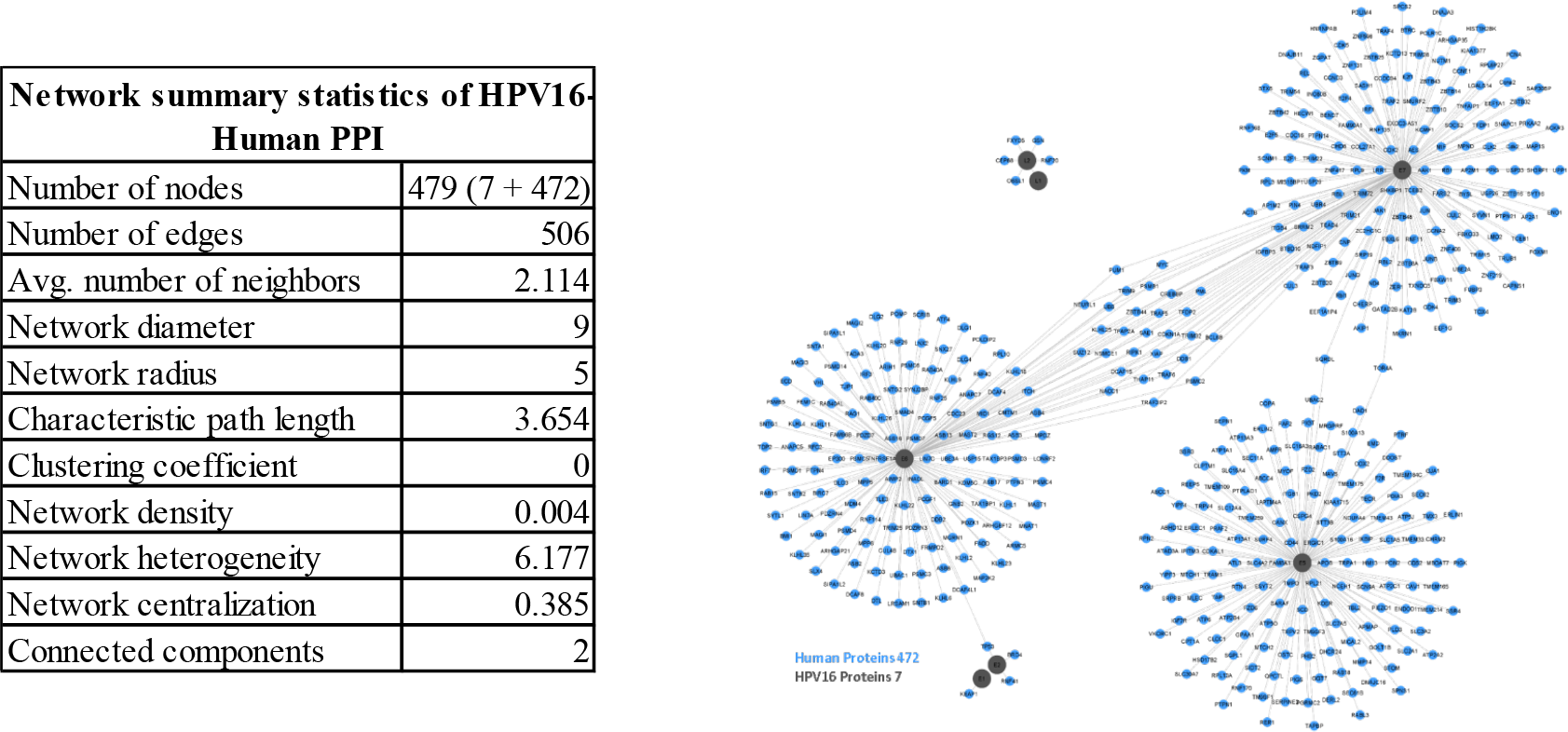
HPV16-Human Protein-Protein Interaction (PPI) Network. 7 HPV viral proteins in dark grey physically interacting with 472 host human proteins in blue. E5, E6, and E7 viral proteins are the hub proteins with the degree of 181, 161, and 152 respectively.

### Enrichment Analysis of Human proteins interacting with HPV16 proteins

After constructing and analysing the viral-host network, we performed an enrichment analysis of the 472 human proteins that interacted with 7 HPV16 viral proteins. Using the EnrichR web interface, we identified 284 significantly enriched GO biological processes, 29 GO molecular functions, 50 GO cellular components, and 76 KEGG pathways (with an adjusted p-value of less than 0.05). Figure 2 shows the top 10 enriched GO biological processes, molecular functions, cellular components, and KEGG pathways. Top KEGG Pathway terms included Epstein-Barr virus infection (p-value: 3e-21), Ubiquitin mediated proteolysis (p-value: 3e-19), and Cell cycle (p-value: 2e-16) with overlapping genes of 36/202, 29/140, and 25/124 respectively. Moreover, top GO Biological Processes included protein ubiquitination (p-value: 2e-38), protein poly-ubiquitination (p-value: 1e-28), and proteasome-mediated ubiquitin-dependent protein catabolic process (p-value: 3e-26) with 77/525, 52/314, and 50/321 respectively. Top GO Molecular Functions included ubiquitin-protein transferase activity (p-value: 8e-34), ubiquitin-protein ligase activity (p-value: 8e-21), and ubiquitin-like protein ligase activity (p-value: 2e-20) with 63/392, 40/263, and 40/270 respectively. Finally, the top GO Cellular Component included cullin-RING ubiquitin ligase complex (p-value: 8e-16), endoplasmic reticulum membrane (p-value: 5e-15), and neuromuscular junction (p-value: 9e-10) with 27/157, 56/712, and 11/38 respectively. These results suggest that the ubiquitin-proteasome system and cell cycle regulation are important processes involved in HPV infection. Further details on the enrichment analysis can be found in Supplementary Table ST4.

**Figure 2:**
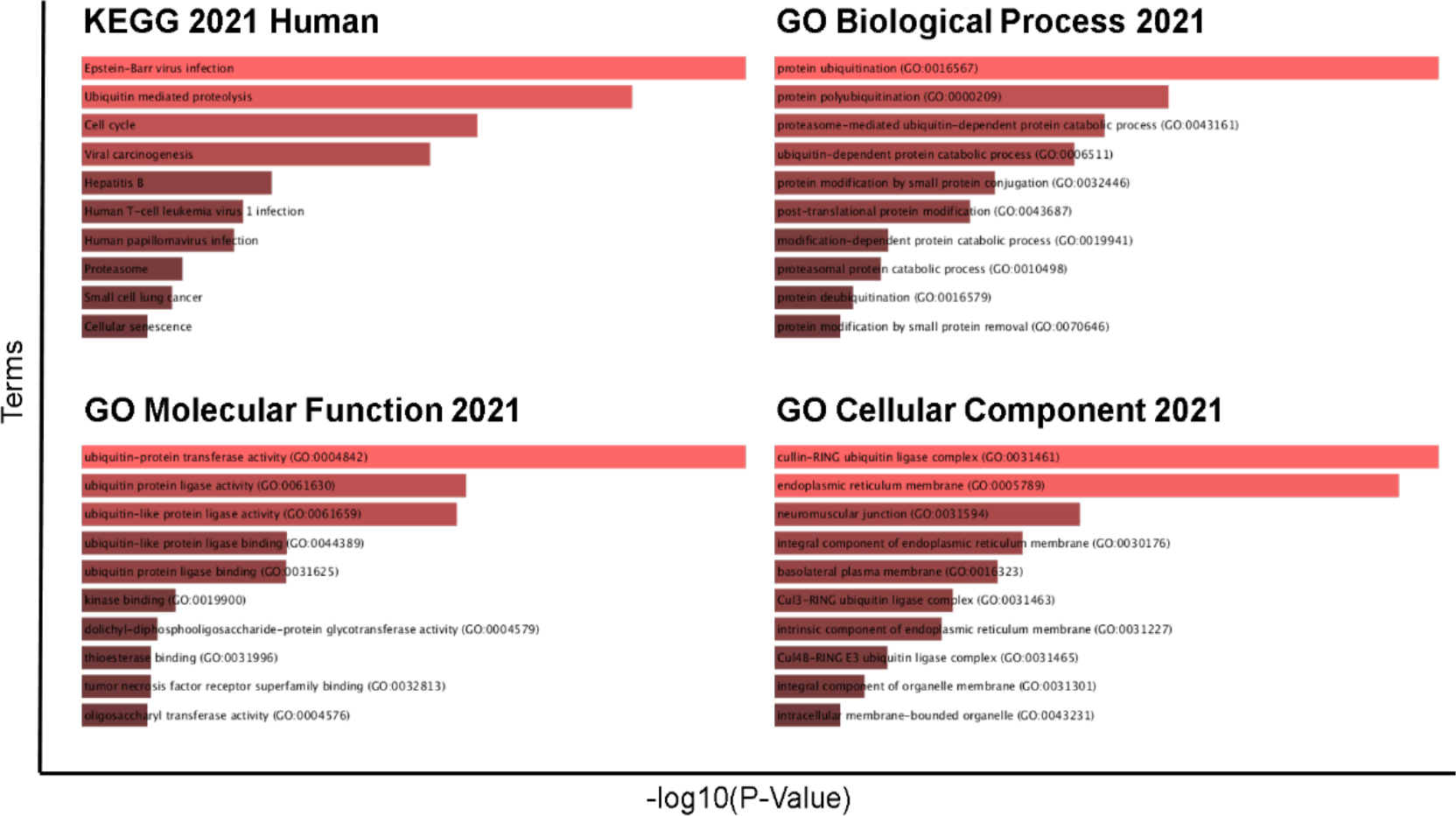
Gene ontology and pathway enrichment analysis of Human Protein interactors of HPV16 proteins. Gene ontology and pathway enrichment analysis of the genes associated with human proteins that interact with HPV16 proteins. The vertical axis represents top 10 GO terms and KEGG pathways while the hortizontal axis represents the statistical significance of the annotations in terms of –log10 (p value).

### HPV16 proteins interact with 37 human proteins involved in OSCC and OPSCC

Following the analysis of the HPV16-human protein-protein interaction (PPI) network, we sought to investigate the relationship between HPV16 and oropharyngeal squamous cell carcinoma (OPSCC) and oral squamous cell carcinoma (OSCC). To achieve this, we compiled a list of proteins that have been implicated in cancer and created a viral-host network specific to OPSCC and OSCC, as illustrated in Figure 3. We obtained a gene list for OPSCC and OSCC based on the frequency of mutations in the oropharynx and oral cavity samples from the TCGA project using data from cBioPortal (see methods). The number of nodes representing human proteins was then reduced from 472 to 37 by filtering out the nodes in the network that were not associated with either OSCC or OPSCC based on the compiled lists. Of these 37 HPV16-OSCC/OPSCC interactors, 19 unique proteins were mapped to OPSCC, 14 to OSCC, and the remaining 4 were common to both cancer types. These findings indicate the existence of a potential mechanistic bifurcation in HPV infection between OSCC and OPSCC (as illustrated in Figure 3).

### HPV16-OSCC/OPSCC interactors show major involvement in cancer-related pathways

After the identification of HPV16-OSCC/OPSCC interactors, we conducted a gene-set enrichment analysis to identify the KEGG pathways and Gene Ontology (GO) terms associated with the set of cancer-related genes whose proteins interact with HPV16. The top 10 enriched terms under Gene Ontology (biological processes, molecular functions and cellular components,) and KEGG pathways for the 37 genes are presented in Figure 4. Among the KEGG pathways, 18 out of 37 genes were linked with Human Papilloma Virus infection, as well as cell cycle and viral carcinogenesis pathways. These pathway terms can provide additional insights into the biological mechanisms of HPV-associated OPSCC. Interestingly, the enrichment analysis of the 37 genes showed significant involvement of the genes in human papillomavirus infections as well as cancer-related pathways and processes (as shown in Figure 4)..

**Figure 4:**
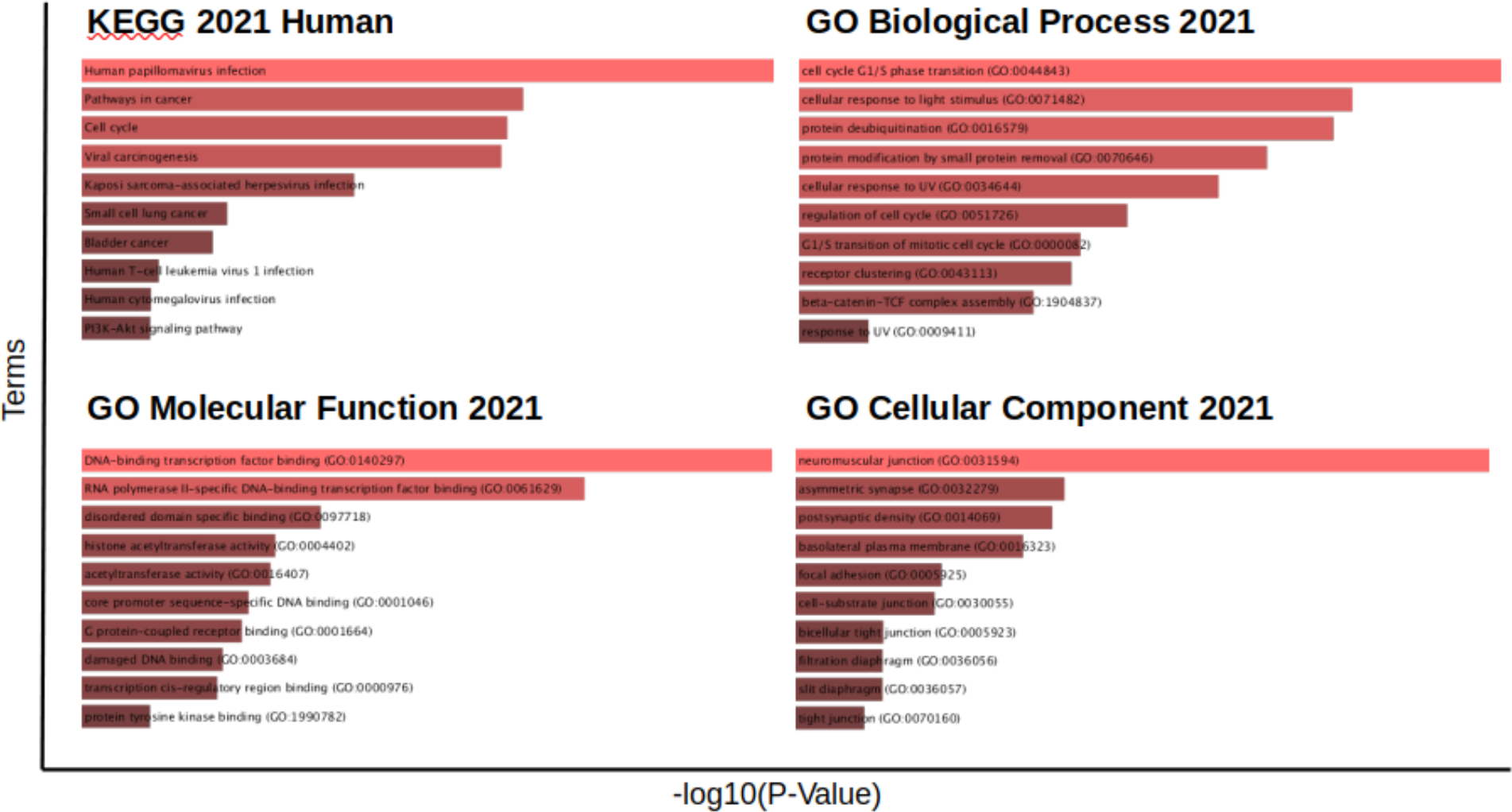
Gene ontology and pathway enrichment analysis of Human Protein interactors of HPV16 proteinsGene ontology and pathway enrichment analysis of the human genes associated with OPSCC and OSCC and their proteins are in interaction with HPV16 proteins. The vertical axis represents top 10 GO terms and KEGG pathways while the horizontal axis represents the statistical significance of the annotations in terms of –log10 (p value).

### NUTM1, MYC, and SCN9A are more differentially expressed in HPV+ OPSCC

We utilized the TCGA dataset to retrieve expression profiles of a similar set of cases used to HPV-OPSCC/OSCC interactors, for which expression profiles were available. After preprocessing the data, we conducted differential gene expression analysis separately for primary tumours of HPV-oral cavity (n=142), HPV+ oral cavity (n=11), HPV-oropharyngeal (n=10), and HPV+ oropharyngeal (n=22), separately against normal head and neck tissue samples (n=31) obtained from the Broad Institute GDAC Firehose. We compiled and filtered the results from the four analyses for 37 genes. As shown in Figure 5, there was no difference in fold change across groups for 22 out of 37 genes, which was expected (volcano plots figure in Supplementary Figures SF1-3).

**Figure 5:**
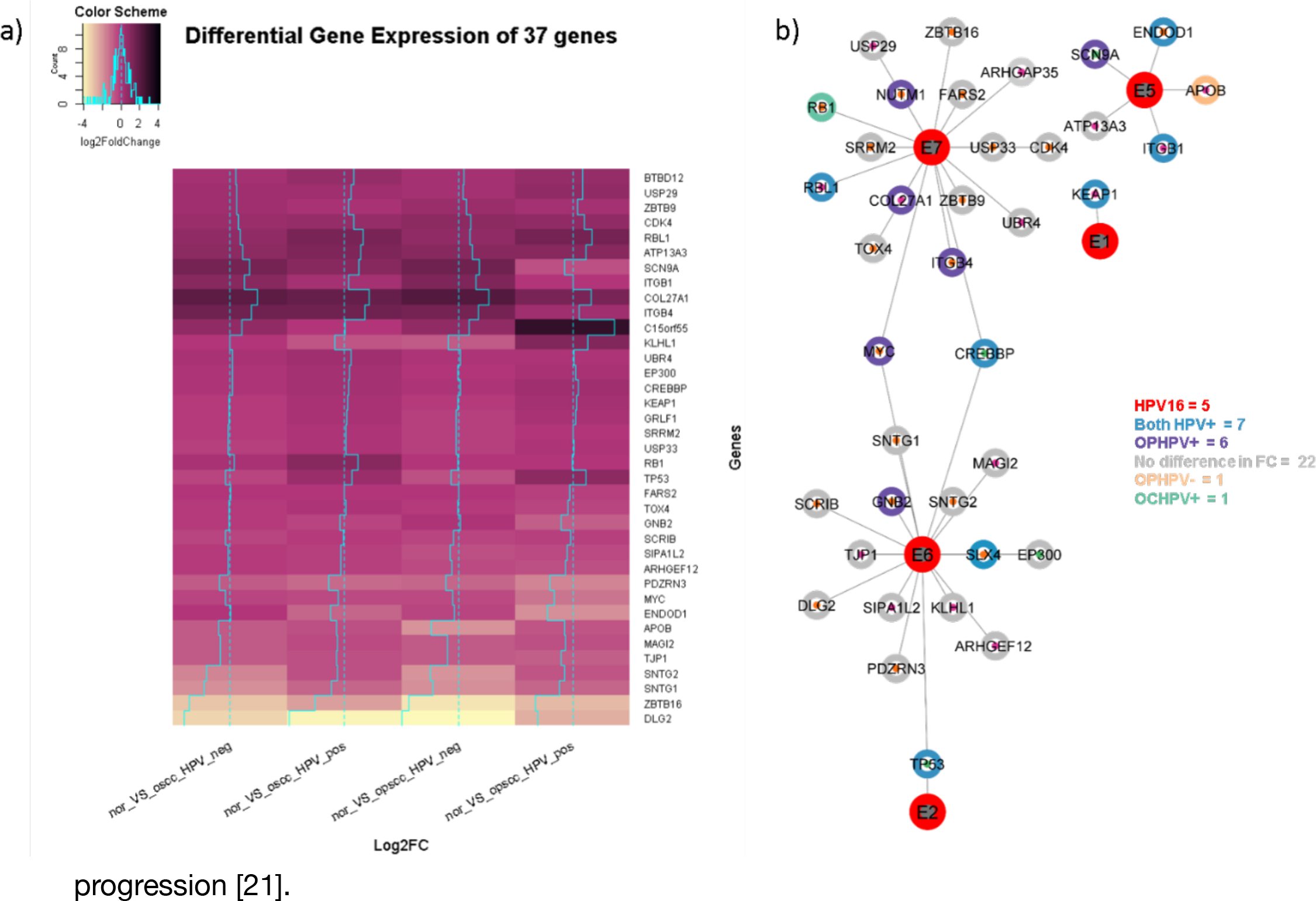
Heatmap of differential gene expression of 37 genes and their connectivity with HPV16 Proteins. (a) Columns represent the fold change of gene expression for experiment OSCC HPV-, OSCC HPV+, OPSCC HPV-, and OPSCC HPV+ relative to the normal, while rows represent individual genes. Black indicates over-expression and light yellow shade indicates under-expression and magenta indicates close to no change in expression. The trace in cyan in each column corresponds to fold change for each gene. (b) Connectivity of 37 proteins with 5 HPV16 proteins. Protein of interest in purple because expression pattern related to OPSCC HPV+. NUTM1 COL27A1, ITGB4 interacts with E7, MYC connects E6 and E7, GNB2 interacts with E6, and SCN9A interacts with E5.

However, there was a difference in expression for 7 genes in both HPV+ OPSCC and HPV+ OSCC. In both HPV-positive cancers, TP53 was over-expressed and ENDOD1 was under-expressed. CREBBP, SLX4 [BTBD12], ITGB1, KEAP1, and RBL1 were slightly under or over-expressed in both HPV+ cancers. RB1 was only over-expressed in OSCC HPV-positive, while APOB was more under-expressed in OPSCC HPV-negative.

An interesting finding from the differential expression analysis was that six genes exhibited a different expression in OPSCC HPV+ compared to the other three groups. NUTM1 [C15orf55] was the most over-expressed in OPSCC HPV+. ITGB4, GNB2, and COL27A1 were slightly under-expressed, while SCN9A and MYC were more under-expressed.

## Discussion

We begin our study by identifying genes that have been found to be mutated in OSCC and OPSCC. For this, we utilized the publicly available data from The Cancer Genome Atlas (TCGA) (Hoadley et al., 2018). Despite skewness in the availability of samples when comparing OPSCC and OSCC, we discovered 612 genes for OSCC and 560 genes for OPSCC (including additional hits from the literature review; refer to Supplementary Table ST3). The potential reason for the lower number of mutated genes in OPSCC is that it is thought to have more viral-driven carcinogenesis than OSCC. During the malignant transformation, the host cell acquires somatic changes due to genome instability caused by E6 and E7, resulting in copy number variations and chromosomal rearrangements [20].

HPV16’s seven proteins i.e. E1, E2, E5, E6, E7, L1, and L2 make 506 physical interactions with 472 human proteins with an average degree of 2.114 (*Table 1*), where E5, E6, and E7 are the hub proteins. Figure 1. When performing KEGG and GO enrichment analysis for the human proteins that are in interactions with the viral proteins, most of the proteins were found to be enriched in ubiquitination (*Figure 2*). which targets protein degradation, signalling, and DNA repair as a result of a mutation or an epigenetic influence that can play a critical role in cancer development and progression [21].

In the context of viral carcinogenesis, ubiquitination can play a pivotal role in the ability of certain viruses to promote the development of cancer. For example, certain viral oncoproteins, such as HPV16’s E6 and E7, can target cellular proteins for ubiquitination and degradation, leading to the disruption of normal cellular processes and the promotion of cancer. Additionally, viruses can also hijack the host cell’s ubiquitination machinery to promote their replication and evade the host’s immune response [22].

We identified that 37 out of 472 human proteins were labeled in the HPV16-Human PPI network after labelling them with genes mutated in OPSCC and OSCC. The OPSCC group contained 19 proteins, while the OSCC group contained 14 proteins, four of which were common. Furthermore, the network in Figure 3 shows that E7 harbours most of the cancerous proteins, with 9 corresponding to OPSCC [ZBTB9, TOX4, SRRM2, USP33, ZBTB16, FARS2, CDK4, ITGB4, RB1] and 5 corresponding to OSCC [COL27A1, USP29, ARHGAP35, UBR4, RBL1]. E6 interacts with 7 OPSCC proteins [PDZRN3, GNB2, SNTG1, SNTG2, DLG2, SCRIB, SLX4] and 5 OSCC proteins [KLHL1, MAGI2, SIPA1L2, TJP1, ARHGEF12]. EP300, on the other hand, is found in both cancer subtypes and interacts with E6. MYC and CREBBP link E6 and E7, while TP53 links E6 and E2. Because E6 and E7 are the most important oncogenic proteins, their influence on OPSCC-labeled proteins is more indicative of the role of HPV16 in OPSCC than OSCC. E5, on the other hand, interacts with three OSCC-labeled proteins [ITGB1, ATP13A3, and APOB], one common [SCN9A], and one OPSCC-labeled protein [ENDOD1], indicating its role in OSCC. Even though E5 is considered a minor oncogenic viral protein, it does play a role in early carcinogenesis by maintaining the host cell’s proliferation potential and avoiding apoptosis [23].

As the study aimed to identify potential genes of interest signatures specific to HPV-induced OPSCC, we performed a comparative differential gene expression analysis between HPV+ OPSCC and OSCC, and HPV-OPSCC and OSCC by building on the mutational landscape in OPSCC and OSCC and their connectedness with the HPV16’s proteins. Based on this analysis, we identified six proteins (NUTM1, MYC, SCN9A, COL27A1, ITGB4, and GNB2) that showed differential expression in HPV-positive OPSCC compared to HPV-positive OSCC, HPV-negative OPSCC, and OSCC.

Although the relationship between these six proteins and HPV infection is not well established as more research is needed to determine the exact role of these genes in HPV-associated OPSCC. However, MYC [24], ITGB4 [25], and GNB2 [26] have been implicated in the PI3K-Akt signalling pathway, which is known to contribute to HPV-associated tumorigenesis [27]. Thus, these proteins are promising candidates for further investigation in the context of HPV-associated OPSCC.

In conclusion, we bifurcated OPSCC and OSCC based on mutational landscape and their connectivity with HPV16 and prioritised six candidates based on comparative differential gene expression in HPV-positive OPSCC. Out of these six candidates MYC, ITGB4, and GNB2 are found to be involved in the PI3K-Akt Signalling pathway, which is crucial for the regulation of basal cells as these are known to be progenitor stem cells and a favourite target of HPV infection. Further, *in vitro* screening of these genes is required for validation.

